# Life history mediates the trade-offs among different components of demographic resilience

**DOI:** 10.1101/2021.06.30.450480

**Authors:** Pol Capdevila, Iain Stott, James Cant, Maria Beger, Gwilym Rowlands, Molly Grace, Roberto Salguero-Gómez

## Abstract

Accelerating rates of biodiversity loss underscore the need to understand how species achieve resilience –their ability to resist and recover from a/biotic disturbances. Yet, the factors determining the resilience of species remain poorly understood, due to disagreements on its definition and the lack of large-scale analyses. Here, we investigate how the life history of 785 natural populations of animals and plants predict their intrinsic ability to be resilient. We show that demographic resilience can be achieved through different combinations of compensation, resistance, and recovery after a disturbance. We demonstrate that these resilience components are highly correlated with life history traits related to the species’ pace of life and reproductive strategy. Species with longer generation times require longer recovery times post-disturbance, while those with greater reproductive capacity have greater resistance and compensation. Our findings highlight the key role of life history traits to understand species resilience, improving our ability to predict how natural populations cope with disturbance regimes.

## Main

Preventing biodiversity loss in the face of global change in a major challenge in ecology and conservation^1,2^. As global change accelerates^3^, species –and the services that they provide^4^– are being lost at an unprecedented rate^5,6^. Still, some species can persist or even increase their abundance despite the increasingly frequent and intense disturbance events consequence of global change^7–9^. Such an ability to persist after a disturbance depends, to a large extent, on the species inherent ability to resist and recover from such events, their resilience^10,11^. Therefore, understanding what makes some species more/less resilient than others is crucial to develop effective management and conservation plans^12^. Yet, the lack of data regarding species’ natural populations responses to disturbances and robust methods to quantify resilience have hampered our understanding of the mechanisms that confer resilience to species^11,13,14^.

Understanding what factors render a species resilient requires knowledge about its population dynamics in the context of disturbances. Recent reviews suggest that studies examining the factors affecting resilience have historically focused on ecological communities or whole ecosystems rather than populations^15,16^. However, these studies operating at high levels of biological organisation often lack the level of detail necessary to identify the underlying processes that modulate species responses to disturbances. In contrast, demographic resilience^10^, *i.e.* the ability of a population to prevail before a disturbance, allows for a nuanced exploration of the mechanisms that confer resilience to natural populations. To quantify demographic resilience, disturbances are defined as external, punctual events that cause changes in population structure^10,17^, *i.e.,* the relative proportion of individuals of different size, ages and/or stages in the life cycle of the population. Such disturbances might lead to a relative over- or under-representation of individuals with high survival and/or reproduction, which ultimately will increase or decrease population size^17,18^. Importantly, demographic resilience is a property of the population, as it depends on the characteristics of its life cycle and the vital rates (survival, development, and reproduction) that shape its persistence^17^. Demographic resilience can be captured by three key components: (1) *resistance*, the ability of a population to avoid a decrease in size after a disturbance; (2) *compensation*, the ability of a population to increase its size after a disturbance; and (3) *recovery time*, the time that a population requires to recover its stable demographic structure after a disturbance event^10^ (Fig. 1). Because these three demographic resilience components are based on vital rates common to any species^19,20^, they can be quantified and compared among species and populations.

**Fig. 1.**
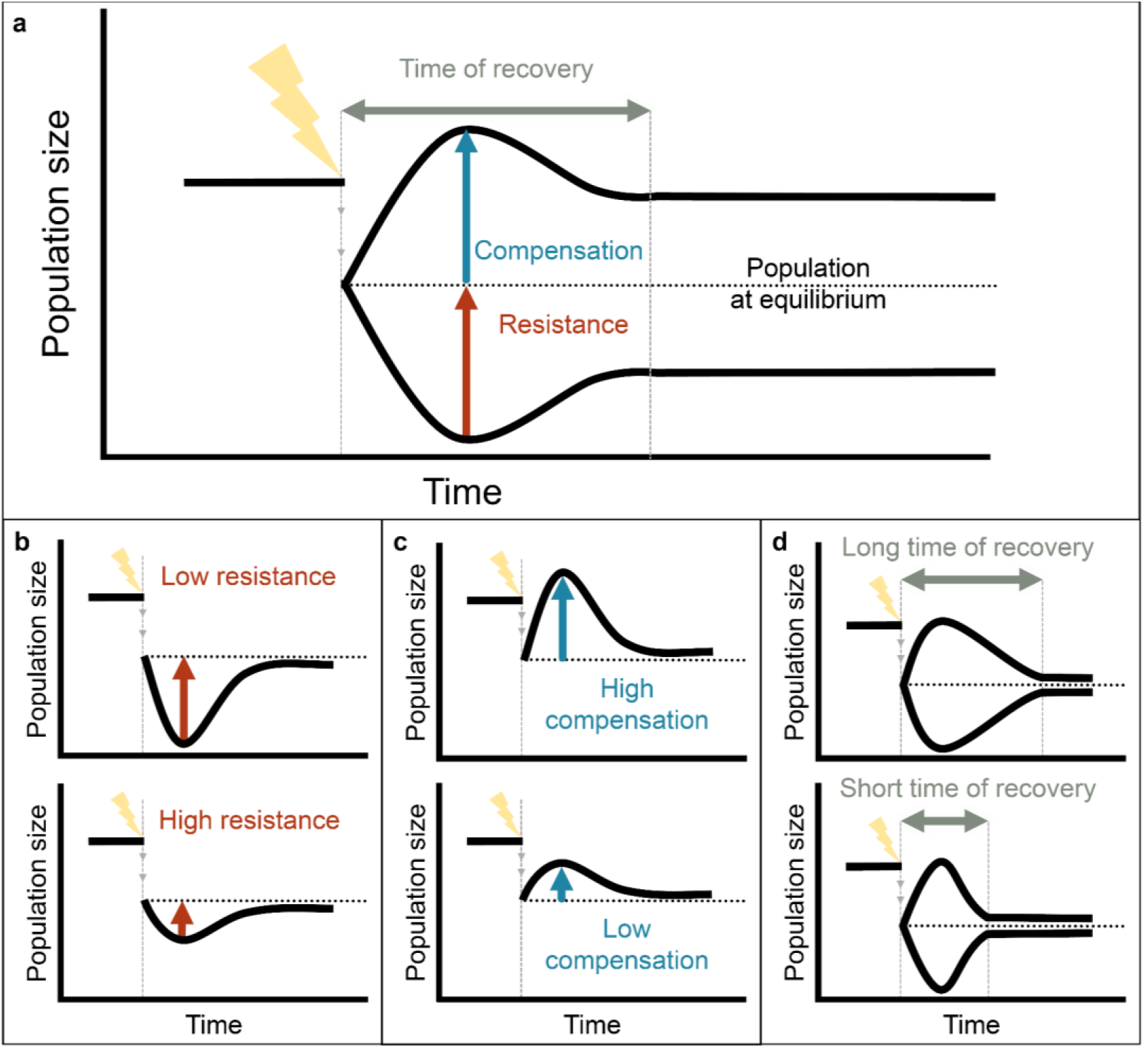
The resilience of a wild population can be quantified via three components: compensation, resistance, and recovery time. **a.** Decomposition of the demographic resilience components of a population affected by a punctual disturbance (lightning bolt). After a disturbance, the size and growth rate of a population may change differently according to how the population structure (*i.e.* the proportion of individuals at different age/stage/size in the population) is affected by the disturbance. **b**. Resistance, the ability to prevent a decline in population size following a disturbance, is measured as the inverse of a population’s decrease following a disturbance relative to its undisturbed conditions (*i.e.* with stable population structure^17^). Low resistance describes high population declines relative to a stable population. **c.** Compensation, the ability to increase relative to the population size following a disturbance, is quantified as the population’s increase with respect to an undisturbed population. Large increases in population size indicate high compensation, while small increases indicate low compensation. **d**. Recovery time, is the period that a population needs to reattain a stable structure after a disturbance.

Determining whether and how the three components of resilience are related to each other is crucial to predict population responses to disturbances. Correlations between resilience components implemented at the community and ecosystem levels often show complex patterns^21,22^. For example, Hillebrand *et al.*^22^, did not find a correlation between resistance and recovery time in experimental plankton communities disturbed by reduced light availability. However, at the species level, resilience components should correlate strongly because they emerge from combinations of vital rates that are under strong selection pressures and frequently trade-off^17^. Naturaly, a significant correlation between two resilience components would allow us to infer one from the other. Importantly, a negative correlation would indicate that the species faces a trade-off when maximising either resilience component. Therefore, understanding the strength and direction of correlations between the components of demographic resilience may help understand and predict whether and how populations may persist despite global change.

The life history strategy of a species is likely a useful proxy to predict its demographic resilience. A species’ life history strategy summarises how limited energy is allocated to survival, development, and reproduction throughout the lifetime of individuals to optimise their fitness^23^. Importantly, life history strategies can influence a species’ response to disturbances^24–28^ and thus its vulnerability to extinction^29–32^. For instance, species with large body size, long generation times, and low reproductive output (*i.e.* slow species, *sensu* Stearns^23^) are often more vulnerable to punctual disturbances than species with small bodies, short generations and highly reproductive^6,33^. As such, “slow” species are expected to be less demographically resilient^34^ to punctual disturbances than “fast” species (*i.e.* specie with small body sizes, short generation times and high reproductive outputs). Still, most of our understanding about the linkages between life histories and demographic resilience comes from theoretical studies using simulated data^35,36^. Besides, the few studies using empirical data have focused on only one or a few components of resilience, such as resistance or recovery^26,28,37^. Consequently, we still lack direct links between the multiple components of resilience (compensation, resistance, and recovery time) and species’ life history strategies.

Here, we examine whether and how the components of demographic resilience trade-off against each other in response to punctual disturbances, and whether they correlate with species’ life history strategies. We use global demographic information for 164 wild populations of 76 animal species and 621 wild populations of 190 plant species (Supplementary Data S1; Table S1), from the open-access databases COMADRE^38^ and COMPADRE^39^, respectively. We also couple these demographic data with phylogenetic information for animals^40^ and plants^41^ separately to account for the lack of independence between the species studied^42^. To establish links between the life history strategy of each species and their demographic resilience, we use these demographic data to estimate key life history traits: generation time (*i.e.*, mean age of reproductive individuals in the population) as an indicator of a species’ pace of life^31^; and mean reproductive output (mean number of recruits produced during the mean life expectancy of an individual in the population) as an indicator of a species’ reproductive strategy^28^. We use hierarchical Bayesian models to test for correlations between the components of demographic resilience, and with species’ life history strategies, and the potential of phylogenetic relationships influencing the observed patterns.

## Results

### Evolutionary history explains more variation in the demographic resilience of animals than in plants

The evolutionary history of the examined species plays an important role in explaining the variation of the demographic resilience of animals, but not in plants (Fig. 2). We assessed the phylogenetic signal fitting hierarchical Bayesian models including the three demographic resilience components as response variable, without fixed effects and including the variance-covariance matrix as the random effect. The explained variance of the random effects was used as a proxy of the phylogenetic signal with a right-skewed distribution close to 0 indicating a lack of phylogenetic signal, and a left-skewed distribution close to 1 suggesting a strong role of phylogenetic relationships in explaining the observed variance (see Methods). The studied animal species show a strong phylogenetic signal for all three components of demographic resilience (Fig. 2**a**), but with a stronger role in the variation of compensation (0.73±0.12, mean±S.E.), and resistance (0.70±0.12) than over recovery time (0.51±0.19). In contrast, evolutionary history plays a minor role in explaining the variation of plant demographic resilience (Fig. 2**b**). In particular, compensation (0.05±0.06) and resistance (0.03±0.05) show a weak phylogenetic signal, compared to recovery time (0.66±0.08). Overall, these results suggest that, while resistance and compensation are highly influenced by the evolutionary history in animals, they are less evolutionarily constrained in plants.

**Fig. 2.**
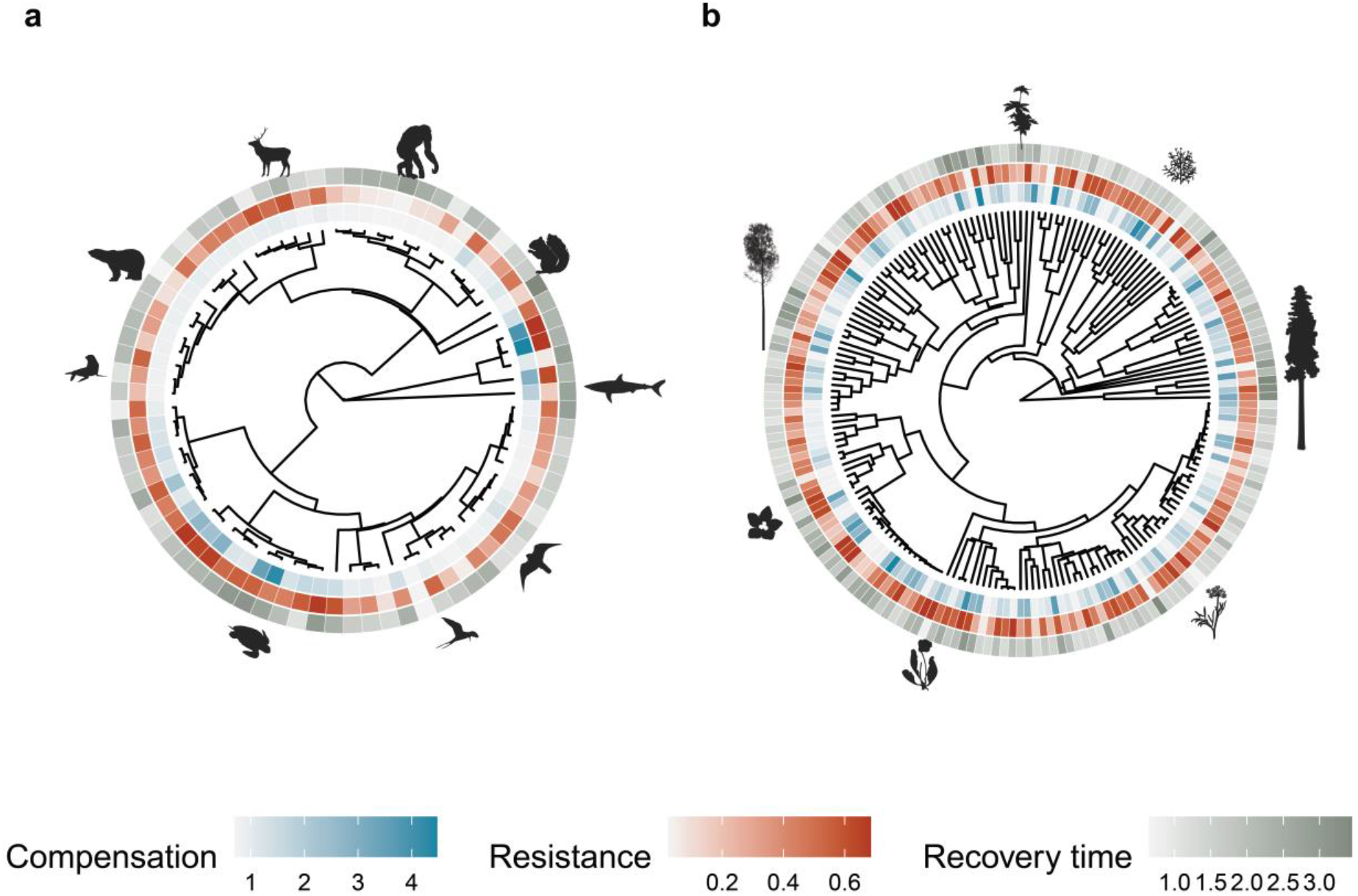
Evolutionary history explains a higher degree of variability of the demographic resilience in animals than in plants. Patterns of variation of demographic compensation, resistance, and recovery time (Fig. 1) for the examined 164 populations of 76 animal species and 621 populations of 190 plant species. Inward ring represents resistance, middle ring compensation, and outer ring recovery time. Evolutionary history explains a greater amount of variability of demographic resilience in animals (**a**) than in plants (**b**). Values showed in each panel represent the mean values of compensation, resistance and recovery time per species. **a**. In animals, the phylogenetic signal was stronger for compensation (0.73±0.12, mean±S.E.), than for resistance (0.70±0.12) and recovery time (0.51±0.19). Silhouettes represent, from the top in clockwise direction, chimpanzee (*Pan troglodytes*), american red squirrel (*Tamiasciurus hudsonicus*), shortfin mako shark (*Isurus oxyrinchus*), peregrine falcon (*Falco peregrinus*), common tern (*Sterna hirundo*), green sea turtle (*Chelonia mydas*), California sea lion (*Zalophus californianus*), polar bear (*Ursus maritimus*), and red deer (*Cervus elaphus*). **b.** In plants, compensation (0.05±0.06) and resistance (0.03±0.05), while recovery time had a stronger phylogenetic signal (0.66±0.08). Silhouettes represent, from the top in clockwise direction, woodland geranium (*Geranium sylvaticum*), wild plantain (*Heliconia acuminata*), white Cypress-pine (*Callitris columellaris*), alpine sea holly (*Eryngium alpinum*), purple pitcher plant (*Sarracenia purpurea*), Douglas’s catchfly (*Silene douglasii*), and grey alder (*Alnus incana*). Silhouettes’ source: phylopic.org.

### Demographic resilience components trade-off strongly to shape species resilience

Our examined species trade-off strongly in demographic components, and so they can achieve demographic resilience either by withstanding disturbances through resistance and compensation, or by minimising recovery time after a disturbance (Fig. 3). We evaluated the association between the components of demographic resilience using the residual correlations from a multivariate hierarchical Bayesian model with compensation, resistance, and recovery time. To account for the lack of independence between the species in our analyses, we used the variance-covariance matrix of phylogenetic distances as a random factor (see Methods). The residual correlations reveal a positive value between resistance and recovery time in animals (Fig. 3**a**). The more resistant to a disturbance an animal is, the longer it needs to recover from the disturbance that pushes its population structure away from its stationary equilibrium. In contrast, resistance and recovery time are negatively correlated in our studied plant species (Fig. 3**a**). Resistance and compensation are also positively correlated in both plants and animals (Fig. 3**b**): species with a high ability to remain stable after a disturbance also have a high ability to compensate after it. Finally, compensation and recovery time are negatively correlated in plants, but positively correlated in animals (Fig. 3**c**).

**Fig. 3.**
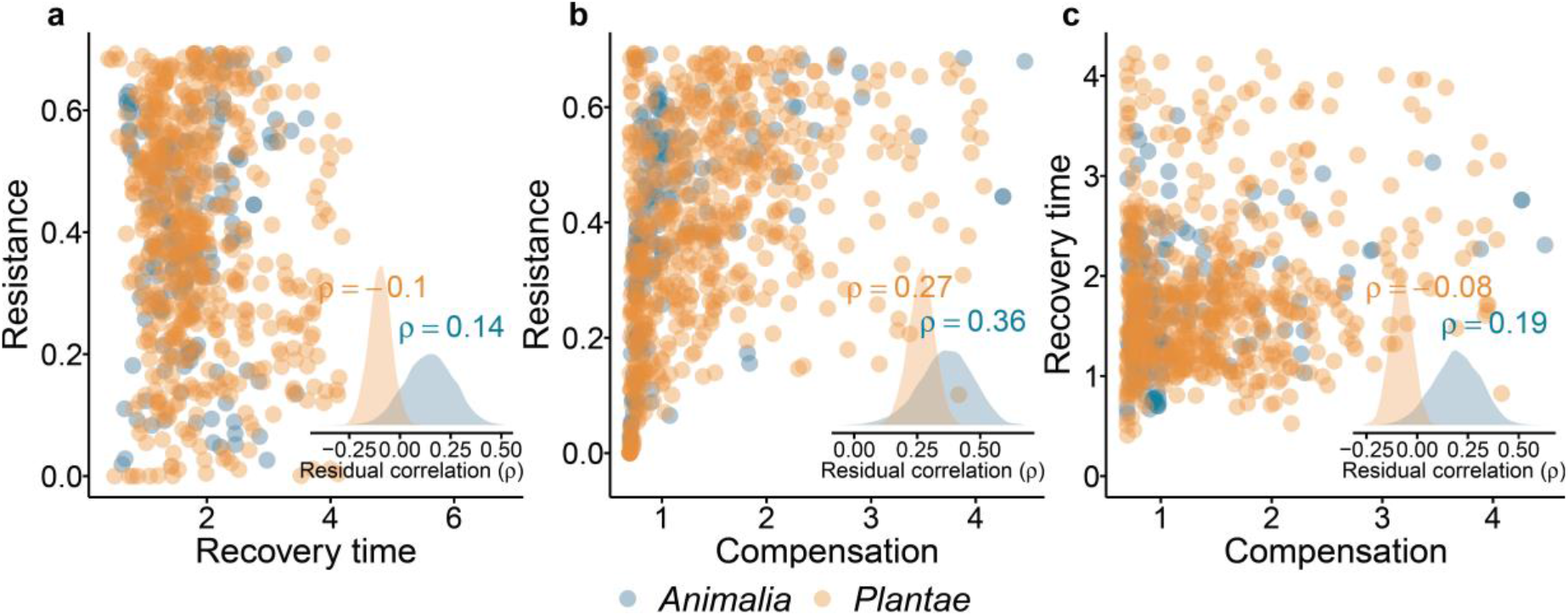
The components of demographic resilience correlate differently for plants than for animals. Correlations between the components of resilience, (**a**) resistance *vs.* recovery time, (**b**) resistance *vs.* compensation, and (**c**) recovery time *vs.* compensation for 164 populations of 76 animal species (blue) and 621 populations of 190 plant species (orange). Insets show the distribution of the residual correlations between the components of resilience, where *ρ* represents the mean value of the distribution. Positive values of *ρ* indicate a positive correlation between components, and negative values represent a trade-off. **a**. The correlation between resistance and recovery time is negative for plants but positive for animals. **b.** Resistance and compensation are positively correlated in both animals and plants. **c**. Recovery time and compensation are slight negatively correlated in plants, but are positively correlated in animals. The residual correlations were estimated by fitting a multivariate hierarchical Bayesian model using compensation, resistance, and recovery time as response variable and with no predictors (see Methods).

### Life history traits predict demographic resilience

The components of demographic resilience are highly correlated with species life history traits (Fig. 4). Generation time is tightly linked to resistance and recovery time, but it has no clear association with compensation (Fig. 4**a-c**; Table S2). However, key differences do occur between plants and animals. While in both groups generation time is positively correlated with recovery time (Fig. 4**c**; Table S2), generation time is negatively correlated with resistance in animals, but positively in plants.

**Fig. 4.**
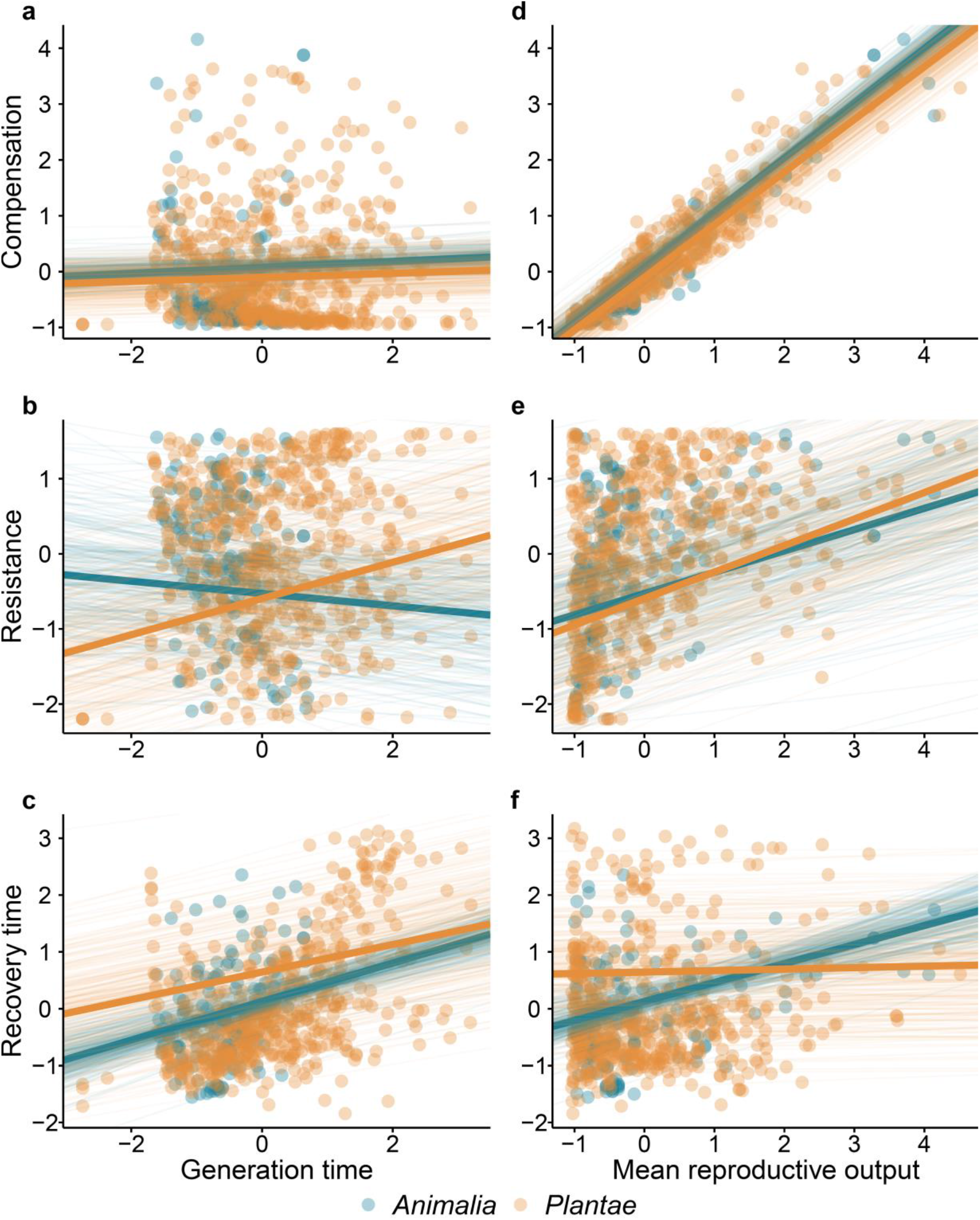
The three components of demographic resilience – resistance, compensation and recovery time – strongly correlate with two key species life history traits: generation time and mean reproductive output. Here we show the correlations between the scaled values of the demographic resilience components of resistance, compensation, and recovery time with the scaled values of generation time and reproductive output of 164 populations of 76 animal species (blue) and 621 populations of 190 plant species (orange). Lines represent the predictions from the hierarchical Bayesian models (Tables S1), where thin lines correspond to the predictions drawn from each of the 250 posterior samples of the model, and the thick line represents the mean outcome of the model.

The mean reproductive output of a species is a strong predictor of its components of demographic resilience (Figure 4**d-f**; Table S2). Here again, though, plants and animals employ slightly contrasting strategies to achieve resilience. Reproductive output is positively correlated with the resistance and compensation of both animals and plants (Figure 4**d-e**; Table S2). However, reproductive output is positively correlated with the recovery time of our studied animals but lacks a clear relationship with the recovery time of plants. Also, our results suggest that being highly reproductive trades-off with longer periods of recovery, at least in the examined animals. The interactive effects of generation time and reproductive output with the components of demographic resilience are consistent across animals and plants, with both kingdoms showing a positive interaction for compensation (Table S2), but a non-significant effect on resistance or recovery time.

## Discussion

We provide empirical support that multicellular species can achieve demographic resilience through different combinations of compensation, resistance, and recovery time. We show that these combinations of demographic resilience are largely determined by the life history strategy of each species. However, the evolutionary history of species plays an important role in animal but not plant demographic resilience. This is perhaps not surprising, since a greater deal of evolutionary plasticity in life history strategies of plants has been previously reported compared to animals^34,43,44^. This plasticity in plant life history trade-offs^34^ could explain the lack of phylogenetic signal in their demographic resilience components. Future studies applying our framework of demographic resilience, while incorporating environmental stochasticity and genetic data, will allow exploring the role of plasticity in the evolution of demographic resilience.

We provide evidence that the components of demographic resilience are correlated, a key finding that contrasts with studies focusing on communities or ecosystems suggesting no direct correlation^16,45,46^. Such a discrepancy likely arises due to the different biological levels at which the components of resilience have been studied up to date^16,46^. Indeed, studies focusing on the physiological responses report a robust correlation between resistance and recovery time^47,48^. Given that our approach is focused on intrinsic population properties^10,17^, each of the demographic resilient components is ultimately the result of a combination of their vital rates^10,18,25^. Supporting this rationale, we show that both compensation and resistance are highly positively correlated with species’ mean reproductive output (Fig. 4). We argue that the strong correlations between the components of demographic resilience that we report here are likely the result of well-known trade-offs between species’ vital rates^23,34^.

The correlation between resistance and recovery time, and between compensation and recovery have opposite but consistent directions in animals and plants. In animals these relationships are positive, whereas in plants they imply trade-offs (negative). Given that compensation and resistance represent population size increase and decrease after disturbance^10,17^, respectively, one would expect populations with a greater ability to compensate (high increase) or resist (lower decline) disturbances to also recover faster. However, it is known that even if populations increase in size quickly after a disturbance or avoid declines in population size, their dynamics will oscillate until they reach their stable structure^17,49^ (structure previous to disturbance in our case). During this transient period, the population has not yet recovered from the disturbance, as the population might experience further increases or decreases in size, which ultimately can lead to local extinction^17,18^. Therefore, even if a population can compensate or resist a disturbance, it might take a long period of time to recover from its effects, no matter how small.

Differences in the relationships among the resilience components in animals and plants have likely emerged from key differences in the evolution of the life history strategies of these two kingdoms. Generation time is linked to the pace of life of species, with longer generation times being associated with slower paces of life^50,51^. However, generation time and mean reproductive output are coordinated differently in plants than in animals. Animals with long generation times usually show low reproductive outputs^51,52^, while in plants (but also some animal groups, such as corals or fishes) species with long generation times can be highly reproductive^34,53,54^. Because there is a tight link between resistance and the mean reproductive output (Fig. 4), it is then not surprising to find divergent relationships between resistance and generation time for both kingdoms.

Beyond generation time and mean reproductive output, there are other life history traits that can influence the reported correlations in demographic resilience components. These traits include animal body weight or plant height, which in turn are highly correlated with other life history traits like generation time and mean reproductive output^34,51^. Using a subsample from our dataset for which body weight and height information is available (93.23% and 43.96% of animal and plant populations, respectively), we observe similar correlations as those for generation time in animals (Fig. S1**a-c**; Table S2). However, these patterns are less clear for plant height (one of the only two currently available proxies for plant body size at a macroecological scale – Fig. S1**a-c**; Table S2) or plant growth form (Fig. S2), another proxy for plant life history strategies^34,55^. These results suggest that having a slow life history strategy results in lower demographic resilience in animals, but not necessarily in plants. Indeed, plants attain alternate resilience strategies depending on their life history strategies: slower plant species do so via greater resistance, while faster plant species attain resilience via shorter recovery times.

To date, much of population ecology and demographic research has focused on how increasing environmentally-driven variability of vital rates (survival, development, and reproduction) impacts the long-term viability of populations^24,27,56^ (but see^25,37,57^). This research has led to the general consensus that species with slow life history strategies tend to be buffered from increased environmental variation^24,27,58,59^. However, this consensus contrasts with the fact that animals with slower paces of life are typically more threatened than species with faster paces of life^29,60^. Although in our analyses we have not found strong links between the conservation status of a species and its demographic resilience (Fig. S3), our results suggest that animals with slow life history strategies often do poorly with disturbances. Our findings suggest that, even if slow paced organisms are buffered against environmental stochasticity^59^, they may still be vulnerable to disturbance events caused by other global change agents.

The resilience of a species is the outcome of multiple factors, and accounting for all of them can be challenging. Our approach evaluates the potential responses of species, based on its intrinsic demographic capabilities^10,17^, to disturbances using the maximum values that a population can increase or decrease after a disturbance. However, the intensity, frequency, duration, and temporal autocorrelation of a disturbance regime can all modify the strength and direction of the relationships among components of resilience in ecological communities^21^. Moreover, since ecosystems are currently exposed to numerous concurrent disturbances^61^, future studies should explore how the interaction between disturbances affects resilience and the correlation among its components^21^. Besides, our approach does not explicitly consider density-dependence, which is known to shape population responses to disturbances^62^. The lack of large volumes of demographic data incorporating density-dependence and analytical tools to properly derive resilience from these data is currently an important barrier to overcoming this limitation.

As global change advances, species are being lost at an unprecedented rate^61,63^, alongside the services that they provide^61^. Here, we show that resilience of natural populations emerges from different combinations of compensation, resistance, and recovery time for animals than for plants. We also demonstrate that life history strategies of species strongly determine their demographic resilience, with important differences among plants and animals. Incorporating knowledge of a species life history is critical to predicting how its populations may respond to disturbances. Understanding how species achieve resilience, as done here, will prove key in developing effective conservation actions.

## Methods

### Data selection

To calculate animal and plant demographic resilience and life history traits we used matrix population models from the COMADRE Animal Matrix Database^38^ and COMPADRE Plant Matrix Database^39^. These databases contain demographic data compiled as age-, size- or developmental stage-structured matrix population models^19^ for over 1,000 plant and animal species. Matrix population models are mathematical representations of a species’ life cycle^19^, where each entry of the projection matrix ****A**** is a product of the vital rates of each stage, size or age of the species. That is, each column of matrix ****A**** contains all contributions by an average individual in a particular class at time *t*, while each row contains all contributions towards the number of individuals in a particular class at time *t*+1:

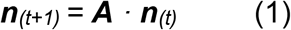

where ****n**** represents the population vector of abundances based on stages, sizes and/or ages. Where the dominant eigenvalue of ****A****, *λ*_1_, represents the asymptotically stable population growth rate^19^, and the right eigenvector ****w**** represents the stable stage structure, that is the relative frequency of individuals in each stage, size and/or age at stationary equilibrium.

To estimate the demographic resilience components and life history traits, we used the individual matrix population models for each available population, after a set of selection criteria (below). When the individual population model ****A**** was not available, we used the mean matrices (e.g. average all ****A**** among different years within a given population). To ensure capturing the natural dynamics of each population, we only included matrices parameterised from non-captive populations in unmanipulated (*i.e.* control) conditions. We also only included described annual dynamics (not seasonal or multiannual) to allow fair comparisons among the different species and populations based on annual time units of population responses to disturbance (e.g. recovery time in years) and life history traits (e.g. generation time in years). To ensure that each matrix population model represented a complete life cycle, we only included those that were irreducible, primitive, and ergodic^64^. The resulting dataset comprised demographic information for 164 populations from 76 species of animals from the COMADRE database^38^, including four populations of Actinopterygii, two Elasmobranchii, 30 Birds, 89 Mammals, and 39 Amphibians (Table S1). The plant data comprised 621 populations from 190 species of plants from COMPADRE database^39^, including 615 populations of Angiosperms and three Gymnosperms (Table S1).

### Demographic resilience

The ****A**** matrix also assumed asymptotic dynamics. That is, these models assumed that the population was at its stable stage distribution ****w****^19^. However, disturbances can change a population’s size and structure, displacing it away from equilibrium structure. Such alterations in population size and structure result in short-term dynamics that can differ from asymptotic dynamics^17^, resulting in either faster or slower growth than that at equilibrium (amplification and attenuation respectively^17^). These transient dynamics represent the intrinsic ability of populations to respond to disturbances, *i.e.* their demographic resilience^10^.

From each selected matrix population model, we estimated three components of resilience: compensation, resistance, and recovery time^10^. Resistance and compensation of each population were studied relative to the long-term population growth or decline predicted by the dominant eigenvalue *λ*_*1*_ of its **A**. To discount the asymptotic effects of the transients and enable fair comparisons among species and populations^17^, we normalised each element of ****A**** by *λ_1_*, resulting in ****Â****, where *λ*_*1*_ = 1 (*i.e.* the population is demographically neither growing nor declining on the long term). Such normalisation avoids the problem of there not being any upper bounds on the transient growth of an asymptotically growing population, or any lower bounds on the transient attenuation of an asymptotically declining population (*sensu* Stott *et al.*^25^). Effectively, this means that all our demographic indices describe how much bigger or smaller a population could get, relative to how big it would be if it started out at stable stage structure^17,25^.

Compensation is estimated as the largest population density that can be reached in the first-time step after disturbance 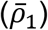, or reactivity^17,18^, which can be calculated as:

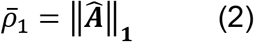

Resistance was estimated as the lowest population density that can be reached in the first time step after disturbance (<u>*ρ*</u>_1_)^17,18^, and was calculated as:

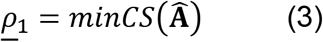

where *minCS* is the minimum column sum of a matrix. eqn 3 values vary from 0 to −1, the more negative the less resistant. To facilitate the interpretation of our resistance estimate, we corrected eqn 3 by adding 1 (**<u>ρ</u>_1_** + 1) so that values closer to 1 correspond to higher resistance.

Recovery time (*t_x_*) was estimated as the time required for the contribution of the dominant eigenvalue (*λ_1_*) to become *x* times as great as that of the subdominant eigenvalue (*λ_2_*), following:

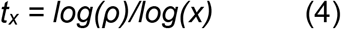

where *x* is set at 10 and *ρ* is estimated as:

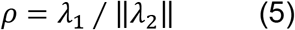

All the transients where calculated using the *popdemo* R package^65^.

### Life history traits and vital rates

From each matrix population model ****A****, we calculated two key life history traits, generation time and mean reproductive output. Generation time represents the mean age of reproductive individuals in the population. Generation time (*T*) was calculated using the function *generation.time* from *popbio* R package^66^, which estimates *T* as:

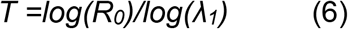

where *R*_*0*_ is the net reproductive rate and *λ*_*1*_ is de dominant eigenvalue of ****A****.

To calculate the mean reproductive output (*φ*), we sum the total sexual reproductive output for each stage of the reproductive component of ****A****, and then weight it by each stage’s relative frequency at stationary equilibrium, as defined by the stable stage distribution ****w****^34^.

### Phylogenetic corrections

To account for the phylogenetic relatedness of the species in our analyses, we used phylogenies for animals and plants. The plant phylogeny was obtained using the *V.PhyloMaker* R package^41^. *V.PhyloMaker* allows to build a rooted and time-calibrated phylogeny using a species list based on already built plant phylogenies^67,68^. The animal phylogeny was produced using the *datelife* R package^69^, a platform that uses publicly accessible phylogenetic source data to build a chronogram – rooted and time-calibrated tree – given an input phylogeny that we sourced from the Open Tree of Life^40,70^. In some cases, for both plant and animal phylogenies, we detected polytomies (*i.e.* >2 species with the same direct ancestor), which can interfere in our phylogenetic signal analyses (see^71^). Polytomies were resolved using the function “multi2di” from *ape* package^72^. Briefly, this approach transforms polytomies into a series of random dichotomies with one or several branches of length very close to 0.

### Statistical analyses

We used different Bayesian hierarchical models with varying structures to address the different research question tackled here. For all the analyses, we log10 transformed resistance, compensation, recovery time, generation time, and mean reproductive output, and z-scaled their values before performing the respective analyses. All the models were fitted using the *brms* package v2.1.0^73^ in R v4.0.0^74^, and run for 8,000 iterations, with a warm-up of 800 iterations. Convergence was assessed visually by examining trace plots and using Rhat values (the ratio of the effective sample size to the overall number of iterations, with values close to one indicating convergence).

To explore the influence of the evolutionary history in determining the patterns of variation of the demographic resilience components, we modelled compensation, resistance, and recovery time across phylogenetic relatedness. We fitted one model per demographic resilience components using hierarchical Bayesian models without fixed effects and including the variance-covariance matrix of the phylogeny and species as a random effects. We assessed the phylogenetic signal by calculating the explained variance of the random effects (phylogeny and population) in the posterior distributions of the models. A distribution pushed up against 0 indicates a lack of phylogenetic signal, given that the variance explained is bounded by 0 and only has positive values. We set weakly informed priors:

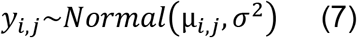

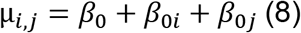

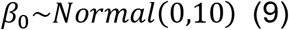

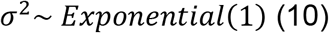

where β_0_ is the global intercept, *β*_0*i*_ and *β*_0*j*_ are the population-level and phylogenetic-level departure from *β*_0_, respectively; *y*_*i*,*j*_ is the estimate for compensation, resistance and recovery time for the *ith* population for the *jth* phylogenetic distance.

To estimate the correlation among the components of demographic resilience, we fitted a multivariate hierarchical model with compensation, resistance, and recovery time as response variables and without predictors^75^. Then, given that there were no predictors in the model, the residual correlations represented the correlation between compensation, resistance, and recovery time. The residual correlation resembles to classical Pearson’s correlations with values varying from −1 to +1, indicating a negative to positive correlations, respectively, and values close to 0 indicating lack of correlation. To account for the lack of independence between the species in our analyses, we used the variance-covariance metric as a random factor (see above). We used a Student’s t-distribution as the likelihood rather than a normal distribution. We used a Student’s t-distribution as the likelihood because this distribution is less sensitive to multivariate outliers^76^. As priors, we used:

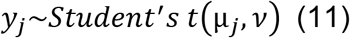

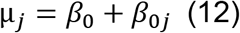

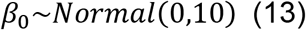

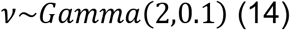

where *β*_0_ is the global intercept, *β*_0*j*_ is the phylogenetic-level departure from *β*_0_; *y*_*j*_ is the estimate for compensation, resistance, and recovery time *jth* phylogenetic distance.

Finally, to estimate the relationship between the components of demographic resilience and the life history strategy of the species, we performed a multivariate hierarchical model using compensation, resistance, and recovery time as response variables and generation time, mean reproductive output, and their interaction as predictors. We also used matrix dimension as a covariate to control for potential confounding effects of the size of the matrix and the resilience component^25^. We also accounted for the lack of independence between the species analysed by incorporating the variance covariance matrix derived from the phylogeny in the model as a random factor. To depict further differences between the studied species not accounted by the phylogeny, we added population as random factor. Because of the different scales of the life history traits and the resilience components, we z-scaled all the variables. For these models, we used weakly regularising normally-distributed priors for the global intercept and slope:

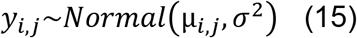

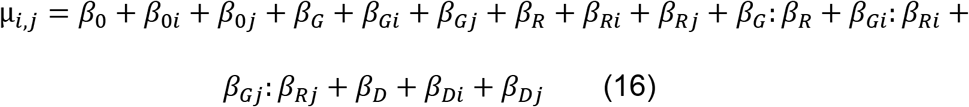

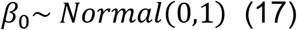

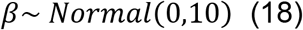

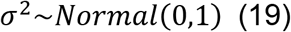

where *β*_0_ is the global intercept, *β*_0*i*_ and *β*_0*j*_ are the population-level and phylogenetic-level departure from *β*_0_, respectively; *y*_*i*,*j*_ is the estimate for compensation, resistance, and recovery time for the *ith* population for the *jth* phylogenetic distance. *β*_*G*_, *β*_*R*_, and *β*_*D*_ represent the effects of generation time, mean reproductive output, and matrix dimension, respectively.

## Acknowledgements

P.C. was supported by a Ramón Areces Foundation Postdoctoral Scholarship (BEVP30P01A5816). RS-G was supported by a NERC IRF (NE/M018458/1). M.G. was supported by a NERC Knowledge Exchange Fellowship (NE/S006125/1). G.R. was supported by an Oxford Martin Fellowship.

## Authors’ contributions

P.C., I. S., M. B. and R.S.-G. conceived the original idea, with contributions from I.S., J.C., G.R. and M.G. P.C., I. S., G. R. and M. G. implemented the statistical analysis. P.C., I.S. and R.S.-G. wrote the initial draft. All authors contributed to the final version of the manuscript.

## Data accessibility statement

– Data and code supporting the results Data will be deposited in the GitHub https://github.com/PolCap/DemographicResilience, upon acceptance of the manuscript. Matrix population models are available at www.compadre-db.org.

## Notes

### Competing Interest Statement

The authors have declared no competing interest.

