## Supplementary for "Life history mediates the trade-offs among different components of demographic resilience"

**Table S1. Taxonomic summary of the matrix population models used in our analyses.** N sp represents the number of species and N pop the number of populations.

| Kingdom | Class | Order | N sp | N pop |
|---|---|---|---|---|
| **Animalia** | Actinopterygii | Perciformes | 3 | 3 |
| | | Siluriformes | 1 | 1 |
| | Elasmobranchii | Lamniformes | 1 | 2 |
| | Aves | Accipitriformes | 5 | 9 |
| | | Anseriformes | 1 | 3 |
| | | Charadriiformes | 3 | 6 |
| | | Falconiformes | 1 | 2 |
| | | Galliformes | 1 | 1 |
| | | Gruiformes | 1 | 1 |
| | | Passeriformes | 1 | 1 |
| | | Pelecaniformes | 1 | 1 |
| | | Procellariiformes | 2 | 2 |
| | | Psittaciformes | 2 | 3 |
| | | Strigiformes | 1 | 1 |
| | Mammalia | Artiodactyla | 5 | 43 |
| | | Carnivora | 12 | 24 |
| | | Chiroptera | 1 | 1 |
| | | Diprotodontia | 1 | 1 |
| | | Primates | 9 | 14 |
| | | Proboscidea | 1 | 1 |
| | | Rodentia | 4 | 5 |
| | Reptilia | Crocodylia | 1 | 1 |
| | | Squamata | 1 | 3 |
| | | Testudines | 8 | 19 |
| **Plantae** | Liliopsida | Alismatales | 1 | 1 |
| | | Asparagales | 5 | 18 |
| | | Dioscoreales | 1 | 1 |
| | | Liliales | 14 | 72 |
| | | Poales | 7 | 19 |
| | | Zingiberales | 2 | 6 |
| | Magnoliopsida | Apiales | 7 | 20 |
| | | Asterales | 20 | 44 |
| | | Brassicales | 10 | 34 |
| | | Caryophyllales | 33 | 84 |
| | | Cornales | 1 | 1 |
| | | Dipsacales | 2 | 6 |
| | | Ericales | 11 | 60 |
| | | Fabales | 12 | 70 |

| | | | | |
|---|---|---|---|---|
| | | Fagales | 4 | 31 |
| | | Gentianales | 3 | 4 |
| | | Geraniales | 2 | 8 |
| | | Lamiales | 11 | 45 |
| | | Magnoliales | 1 | 1 |
| | | Malpighiales | 10 | 23 |
| | | Malvales | 4 | 8 |
| | | Myrtales | 3 | 12 |
| | | Proteales | 1 | 2 |
| | | Ranunculales | 11 | 31 |
| | | Rosales | 4 | 7 |
| | | Sapindales | 4 | 4 |
| | | Saxifragales | 1 | 1 |
| | | Solanales | 2 | 2 |
| | Pinopsida | Pinales | 4 | 6 |

18

**Table S2. Model outputs for the correlations among the components of demographic resilience: compensation, resistance, and recovery time.** Median represents the median of the posterior distribution. CI low and high are the lower and higher values of the 95% confidence intervals, respectively. Rhat is the ratio of the effective sample size to the overall number of iterations, with values close to one indicating convergence values.

| Kingdom | Response | Parameter | Median | CI_low | CI_high | Rhat |
|---|---|---|---|---|---|---|
| **Animals** | Compensation | Intercept | 0.01 | -0.09 | 0.12 | 1.00 |
| | | Generation time | 0.08 | 0.04 | 0.13 | 1.00 |
| | | Mean reproductive output | 0.99 | 0.94 | 1.04 | 1.00 |
| | | Generation time: Mean reproductive output | 0.06 | 0.02 | 0.10 | 1.00 |
| | | Matrix dimension | 0.00 | -0.05 | 0.05 | 1.00 |
| | Resistance | Intercept | -0.42 | -1.73 | 0.91 | 1.00 |
| | | Generation time | -0.12 | -0.28 | 0.04 | 1.00 |
| | | Mean reproductive output | 0.29 | 0.10 | 0.48 | 1.00 |
| | | Generation time: Mean reproductive output | -0.06 | -0.16 | 0.04 | 1.00 |
| | | Matrix dimension | 0.15 | 0.02 | 0.29 | 1.00 |
| | Recovery time | Intercept | 0.14 | -0.14 | 0.62 | 1.00 |
| | | Generation time | 0.31 | 0.20 | 0.42 | 1.00 |
| | | Mean reproductive output | 0.33 | 0.21 | 0.46 | 1.00 |
| | | Generation time: Mean reproductive output | -0.07 | -0.14 | 0.01 | 1.00 |
| | | Matrix dimension | 0.49 | 0.37 | 0.60 | 1.00 |
| **Plants** | Compensation | Intercept | -0.08 | -0.63 | 0.49 | 1.00 |
| | | Generation time | 0.04 | 0.00 | 0.07 | 1.00 |
| | | Mean reproductive output | 0.94 | 0.91 | 0.97 | 1.00 |
| | | Generation time: Mean reproductive output | 0.05 | 0.03 | 0.08 | 1.00 |
| | | Matrix dimension | 0.15 | 0.10 | 0.19 | 1.00 |
| | Resistance | Intercept | -0.64 | -2.12 | 0.80 | 1.00 |
| | | Generation time | 0.24 | 0.14 | 0.34 | 1.00 |
| | | Mean reproductive output | 0.36 | 0.28 | 0.43 | 1.00 |
| | | Generation time: Mean reproductive output | 0.03 | -0.05 | 0.09 | 1.00 |
| | | Matrix dimension | 0.15 | 0.03 | 0.27 | 1.00 |
| | Recovery time | Intercept | 0.60 | -0.89 | 2.13 | 1.00 |

| | | | | | | |
|---|---|---|---|---|---|---|
| | | Generation time | 0.24 | 0.17 | 0.32 | 1.00 |
| | | Mean reproductive output | 0.03 | -0.03 | 0.09 | 1.00 |
| | | Generation time: Mean reproductive output | -0.05 | -0.11 | 0.00 | 1.00 |
| | | Matrix dimension | 0.23 | 0.12 | 0.34 | 1.00 |

*Effects of body size on demographic resilience*

To test the influence of the body size of a species on its demographic resilience, we fitted separated multivariate multilevel Bayesian model for animals and plants. We used compensation, resistance, and recovery time as response variables, and adult body weight (g) for animals or maximum height (m) in plants as fixed effects. We obtained adult body mass (g) data from Myhrvold *et al.*[1] for mammals birds, reptiles and amphibians, and from FishBase[2] for the only elasmobranch species in the study. For terrestrial plants, we utilised maximum height (m) reported per species in the TRY database[3], complemented with information from the Botanical Information and Ecology Network[4] (BIEN; http://bien.nceas.ucsb.edu/bien/). Not all our species had body dimension information available from these databases or via an online search, and so this limitation reduced our initial sample size. The number of populations of animals decreased from 164 to 153, while for plants it decreased from 621 to 273 populations.

For the models, we used weakly regularising normally-distributed priors for the global intercept and slope:

$$y_{i,j} \sim Normal(\mu_{i,j}, \sigma^2) \quad (15)$$

$$\mu_{i,j} = \beta_0 + \beta_{0i} + \beta_{0j} + \beta_B + \beta_{Bi} + \beta_{Bj} + \beta_D + \beta_{Di} + \beta_{Dj} \quad (16)$$

$$\beta_0 \sim Normal(0,1) \quad (17)$$

$$\beta \sim Normal(0,10) \quad (18)$$

$$\sigma^2 \sim Normal(0,1) \quad (19)$$

where $\beta_0$ is the global intercept, $\beta_{0i}$ and $\beta_{0j}$ are the population-level and phylogenetic-level departure from $\beta_0$, respectively; $y_{i,j}$ is the estimate for compensation, resistance and recovery time for the *ith* population for the *jth* phylogenetic distance. $\beta_B$ and $\beta_D$ represent the effects of the animal adult body size (g) or plant maximum height (m) and matrix dimension, respectively.

Body dimension influences the components of demographic resilience of species. Both for animals and plants, as the body size increases, compensation abilities increase (Fig. S1 **a,d**; Table S2). While this pattern would be expected in plants, where large individuals tend to be highly reproductive and then have a greater ability to compensate mortality events[5,6], in most animals larger body sizes are linked to lower reproductive values[7]. Resistance is independent from body dimension for both animals and plants, with the slopes of these correlations showing no clear trend (Fig. S1**b,e**; Table S2). Finally, animal body size is positively correlated with recovery time (Fig. S1**c**; Table S2), while plant body height only shows a slight positive trend (Fig. S1**f**; Table S2).

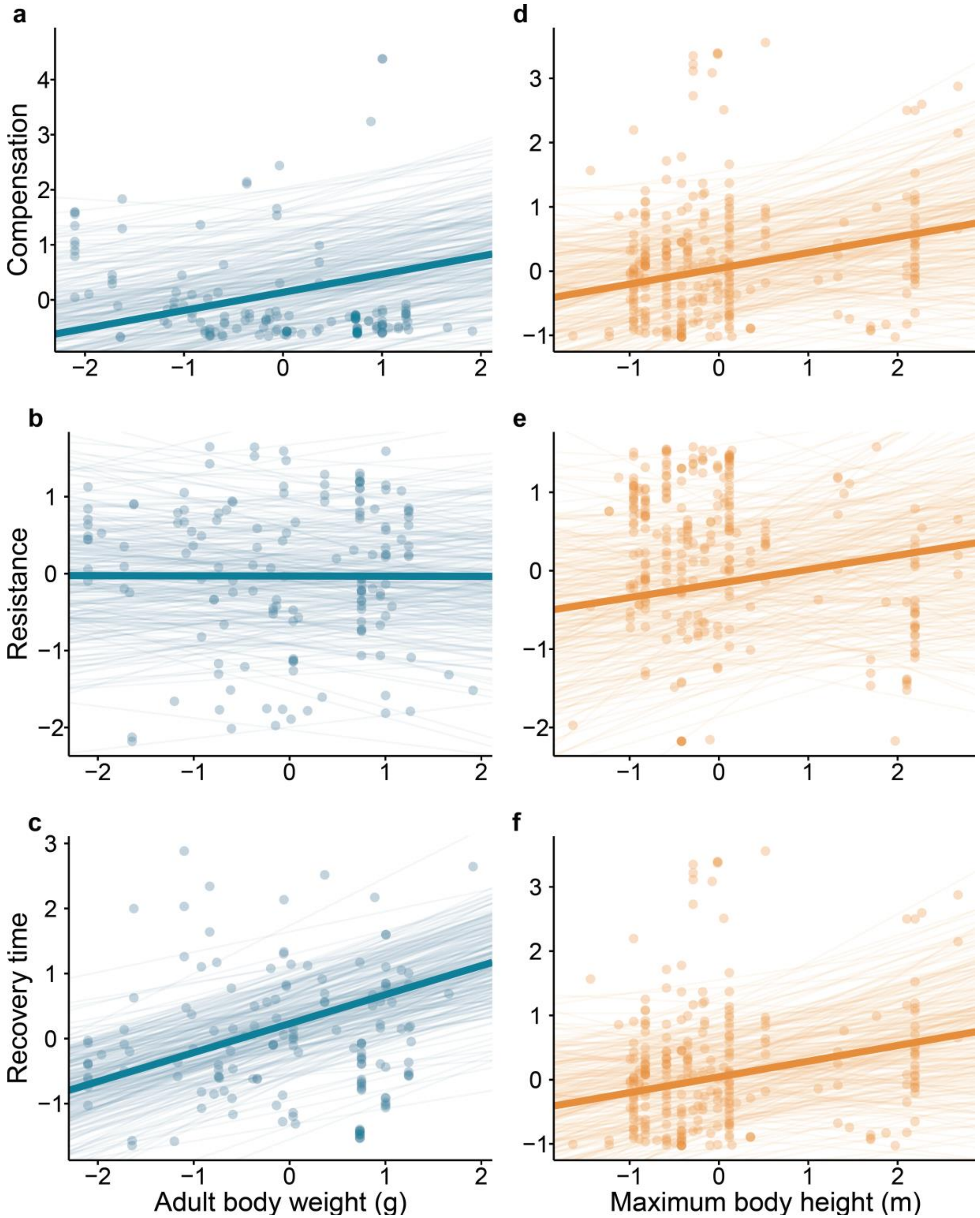


**Fig. S1. Correlation between the components of demographic resilience and body dimensions of animals (a, b, and c) and plants (d, e, and f).** The correlations between the scaled values of the demographic resilience components of resistance, compensation, and recovery time with the scaled values of adult body weight (g) of 149 populations of animals (blue) and 254 plants (orange). Lines represent the predictions from the multivariate multilevel Bayesian models (Table S2), where thin lines correspond to the predictions drawn from each of the 250 posterior samples of the model, and the thick line represents the mean outcome of the model.

**Table S2. Model outputs for the correlations between the components of resilience and body dimension for animals and plants.** Median represents the median of the posterior distribution. CI low and high are the lower and higher values of the 95% confidence interval. Rhat is the ratio of the effective sample size to the overall number of iterations, with values close to one indicating convergence values.

| Kingdom | Response | Parameter | Median | CI_low | CI_high | Rhat |
|---|---|---|---|---|---|---|
| **Animals** | Compensation | Intercept | 0.08 | -1.43 | 1.65 | 1.00 |
| | | Body mass | 0.34 | 0.02 | 0.65 | 1.00 |
| | | Matrix dimension | -0.14 | -0.24 | -0.05 | 1.00 |
| | Resistance | Intercept | -0.08 | -1.42 | 1.18 | 1.00 |
| | | Body mass | 0.00 | -0.30 | 0.31 | 1.00 |
| | | Matrix dimension | 0.09 | -0.06 | 0.23 | 1.00 |
| | Recovery time | Intercept | 0.26 | -0.86 | 1.19 | 1.00 |
| | | Body mass | 0.45 | 0.21 | 0.67 | 1.00 |
| | | Matrix dimension | 0.47 | 0.33 | 0.59 | 1.00 |
| **Plants** | Compensation | Intercept | 0.08 | -1.13 | 1.40 | 1.00 |
| | | Body height | 0.24 | -0.18 | 0.66 | 1.00 |
| | | Matrix dimension | 0.16 | -0.06 | 0.37 | 1.00 |
| | Resistance | Intercept | -0.18 | -1.50 | 1.27 | 1.00 |
| | | Body height | 0.16 | -0.28 | 0.64 | 1.00 |
| | | Matrix dimension | -0.02 | -0.26 | 0.19 | 1.00 |
| | Recovery time | Intercept | 0.16 | -1.10 | 1.40 | 1.00 |
| | | Body height | 0.28 | -0.08 | 0.66 | 1.00 |
| | | Matrix dimension | 0.23 | 0.08 | 0.38 | 1.00 |

*Effects of plant growth form on demographic resilience*

To test whether plant growth form shapes its demographic resilience, we fitted a multivariate multilevel Bayesian model. We used compensation, resistance, and recovery time as response variables and the life form categorisation of Raunkiær[8] as a fixed effect. We classified each plant species according to Raunkiær's growth form scheme[8], in which the height of the shoot apical meristems (SAMs) in relation to ground level determines their life form. Briefly, the levels used here include: (i) helophyte: SAMs under water; (ii) geophyte: SAMs underground; (iii) hemicryptophyte: SAMs at ground level; (iv) chamaephyte: SAMs 0-0.25 m height; (v) nanophanerophyte: SAMs 0.25-8 m; (vi) mesophanerophyte: SAMs 8-30 m; and (vii) megaphanerophyte: SAMs >30m. Importantly, Raunkiær's growth form is often associated with the size or the plant, but also accounts for the shape of the plant and its ability to fully regenerate[8]. Due to lack of information online for some species, our initial sample size from 621 populations was constrained to 437 populations in this specific test.

We used weakly regularising normally-distributed priors for the global intercept and slope:

$$y_{i,j} \sim Normal\left(\mu_{i,j}, \sigma^2\right) \quad (15)$$

$$\mu_{i,j} = \beta_0 + \beta_{0i} + \beta_{0j} + \beta_R + \beta_{Ri} + \beta_{Rj} + \beta_D + \beta_{Di} + \beta_{Dj} \qquad (16)$$

$$\beta_0 \sim Normal(0,1) \quad (17)$$

$$\beta \sim Normal(0,10) \quad (18)$$

$$\sigma^2 \sim Normal(0,1) \quad (19)$$

where $\beta_0$ is the global intercept, $\beta_{0i}$ and $\beta_{0j}$ are the population-level and phylogenetic-level departure from $\beta_0$, respectively; $y_{i,j}$ is the estimate for compensation, resistance and recovery time for the *ith* population for the *jth* phylogenetic distance. $\beta_R$ and $\beta_D$ represent the effects of the Raunkiær life form and matrix dimension, respectively.

Our examined plant species have different values of compensation, resistance and recovery time depending on their Raunkiær life form[8]. Still, the overlap among the posterior estimates is considerable (Fig S2), so that species with different growth forms can show similar levels of demographic resilience. Compensation increased from Helophytes to Macrophanerophytes (Fig. S2), indicating that plant species whose shot apical meristems overwinter at higher distance from the ground have a higher ability to compensate from disturbances. On the contrary, resistance shows no clear patterns across Raunkiær life forms (Fig. S2), suggesting that resistance is independent of the location of SAMs on the plant anatomy. Finally, recovery time displays a concave shape (Fig. S2) from Helophytes to Macrophanerophytes, indicating that plants with SAMs very close to or far from the ground level have the shortest recovery times post disturbance. These results are in agreement with previous studies[5,6], were trees were reported to have a greater ability to bounce back from disturbances than other plant growth forms, due to their higher reproductive outputs. However, we also note a high degree of overlap in the various index of demographic resilience across life forms.

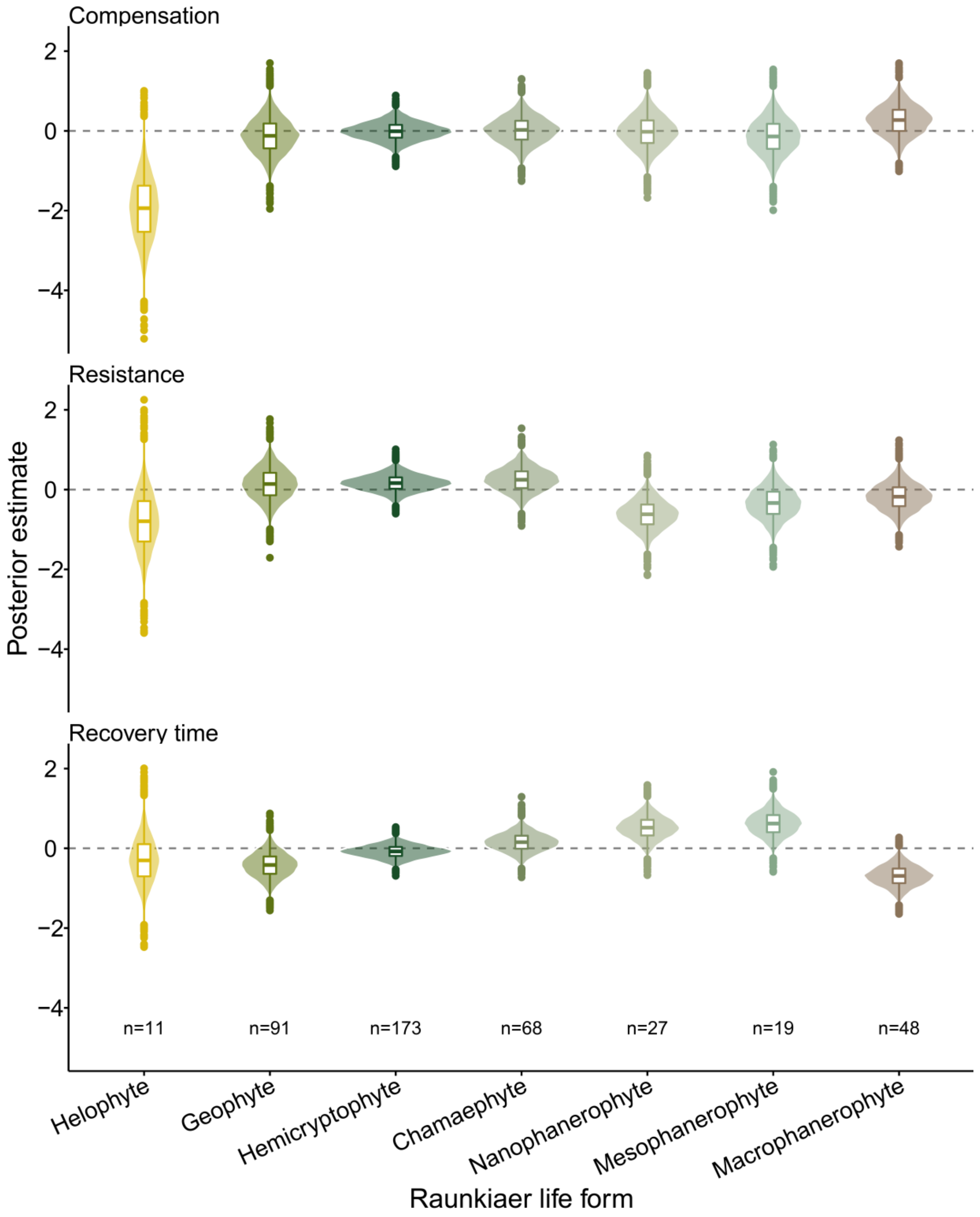


**Fig. S2. Demographic resilience components of plants classified according to Raunkiær life forms.** We show here the posterior distributions of multivariate multilevel Bayesian models with scaled values of compensation, resistance and recovery time as response variable and the Raunkiaer life form classification[8]. n= indicates the sample size of the original dataset.

*Conservation status*

To test whether there was any correlation between the demographic resilience of the species and their conservation status, we fitted two separate multivariate multilevel Bayesian models for animals and plants. We used compensation, resistance and recovery time as response variables and the conservation status and the conservation status of the studied species as reported on the IUCN Red List[9] as fixed effects. Not all the included species had a conservation status available in the IUCN Red List, reducing our initial sample size from 164 to 155 populations of animals, and from 621 to 212 populations of plants.

For these models we used weakly regularising normally-distributed priors for the global intercept and slope:

$$y_{i,j} \sim Normal(\mu_{i,j}, \sigma^2) \quad (15)$$

$$\mu_{i,j} = \beta_0 + \beta_{0i} + \beta_{0j} + \beta_C + \beta_{Ci} + \beta_{Cj} + \beta_D + \beta_{Di} + \beta_{Dj} \quad (16)$$

$$\beta_0 \sim Normal(0,1) \quad (17)$$

$$\beta \sim Normal(0,10) \quad (18)$$

$$\sigma^2 \sim Normal(0,1) \quad (19)$$

where $\beta_0$ is the global intercept, $\beta_{0i}$ and $\beta_{0j}$ are the population-level and phylogenetic-level departure from $\beta_0$, respectively; $y_{i,j}$ is the estimate for compensation, resistance and recovery time for the *ith* population for the *jth* phylogenetic distance. $\beta_C$, and $\beta_D$ represent the effects of the conservation status and matrix dimension, respectively.

There was a high variation in the compensation, resistance and recovery time among the different conservation statuses of the species (Fig. S3). Interestingly, these results suggest the current conservation status of species is independent of their inherent ability to compensate, resist and recover from disturbances.

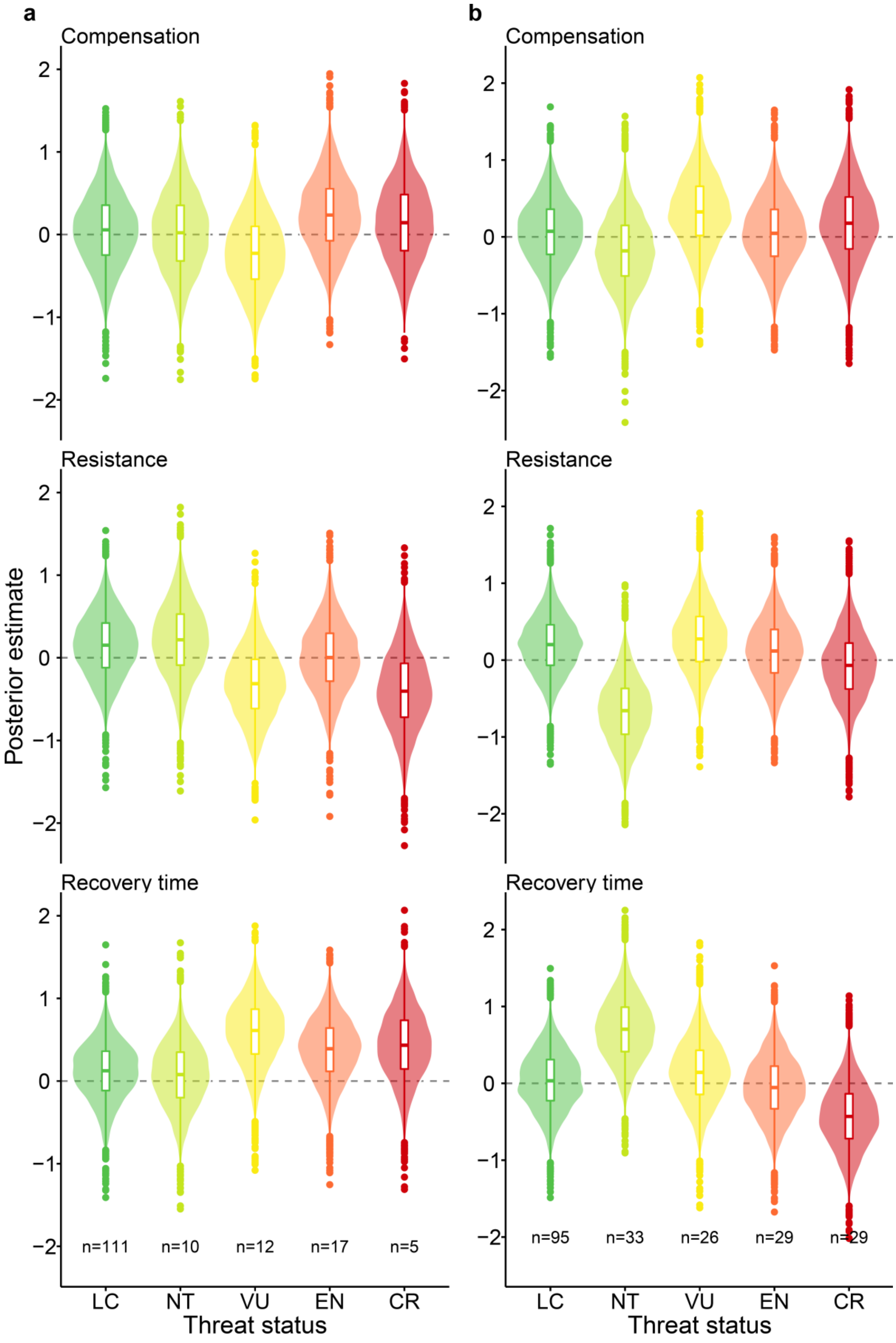

a
Compensation
Resistance
Recovery time
Posterior estimate
n=111
n=10
n=12
n=17
n=5
LC
NT
VU
EN
CR
Threat status
b
Compensation
Resistance
Recovery time
n=95
n=33
n=26
n=29
n=29
LC
NT
VU
EN
CR
Threat status


155

**Fig. S3. Demographic resilience components of species against their conservation status in animals (a) and plants (b).** We show here the posterior distributions of multivariate multilevel Bayesian models for animals (**a**) and plants (**b**), with scaled values of compensation, resistance and recovery time as response variable and the conservation status of the studied species as reported on the IUCN Red List[9] as fixed effects. LC = Least Concerned, NT = Near Threatened, VU = Vulnerable, EN = Endangered, CR = Critically Endangered. n = indicates the sample size.
